# Distinct brainstem to spinal cord noradrenergic pathways differentially regulate spinal neuronal activity

**DOI:** 10.1101/2021.09.25.461801

**Authors:** Mateusz Wojciech Kucharczyk, Francesca Di Domenico, Kirsty Bannister

**Author notes:** **Corresponding author:** Address: Institute of Psychiatry, Psychology and Neuroscience, Wolfson CARD, Guy’s Campus, King’s College London, London, SE1 1UL. UK. **Senior author** Address: Institute of Psychiatry, Psychology and Neuroscience, Wolfson CARD, Guy’s Campus, King’s College London, London, SE1 1UL. UK.

## Abstract

Brainstem to spinal cord pathways modulate spinal neuronal activity. We implemented locus coeruleus (LC) targeting strategies by microinjecting CAV-PRS-ChR2 virus in the spinal cord (LC:SC module) or LC (LC:LC module). While activation of both modules inhibited evoked spinal neuronal firing via α_1_-adrenoceptor-mediated actions, LC:SC opto-activation abolished diffuse noxious inhibitory controls. The LC as a pain generator is likely mechanistically underpinned by maladaptive communication with discrete descending modulatory pathways.

## Main

The descending pain modulatory system (DPMS) comprises noradrenergic projections that underpin a tonic pathway from the locus coeruleus (LC) to the dorsal horn of the spinal cord^1^ and diffuse noxious inhibitory controls (DNIC). Broadly, LC and DNIC pathways represent distinct forms of spinal adrenoceptor (AR) driven endogenous analgesia^2–4^. However, recent evidence of a ‘potentiated inhibitory' effect on pain-related behaviours following antagonism of spinal α_2_-ARs^5^ and opposing α_2_-AR-mediated facilitatory signalling in the brainstem^6^ highlights the complexity of the pharmacological functionality of the pathways therein. This complexity extends to the transition from acute to chronic pain when considering the modular functional organisation and ‘chronic pain generator' role of the LC^7,8^, as well as disease-stage specific DNIC expression in rodent models of chronic pain^9,10^.

Previously, chemogenetic activation of descending noradrenergic controls following microinjection of spinal CAV-PRS virus inhibited wide dynamic range (WDR) neuronal activity. This inhibition was reversed by spinal atipamezole (α_2_-AR antagonist)^7^. Conversely, optoactivation of the LC following LC CAV-PRS microinjection (thus labelling a LC:LC module) caused inhibition of the spinal reflex only with ventral optic fibre placement, whereupon atipamezole no longer reversed the inhibitory effect^5^. Hypothesising that this differential atipamezole impact reflected the abolished action of a non-coerulean inhibitory noradrenergic control, namely DNIC, we microinjected CAV-PRS-ChR2 spinally (thus labelling a LC:SC module) or LC proper (LC:LC module) (Fig. 1a). Following confirmation of ipsilateral ventral LC labelling (ventral: 12.4±2.4% vs. dorsal: 4.0±1.5% ipsilateral noradrenergic LC neurons expressed mCherry) (Fig. 1b), it was demonstrated that optoactivation of both LC:LC and LC:SC modules (Fig. 1c, S1) inhibited mechanically-evoked spinal WDR neuron activity (Fig. 1d-j). This inhibition was reversed by α_1_-AR antagonist prazosin (Fig. 1k-l) but enhanced by local application of atipamezole (Fig. 1m-n). These results suggest that activation of the LC inhibits spinal WDR neuron activity via a α_1_-AR-mediated mechanism. Further, the results imply that the impact of atipamezole on WDR activity in the study by Hirschberg and colleagues^7^ was likely a result of the malfunction of another, non-coerulean, inhibitory noradrenergic control. Proposing DNIC to be the affected circuit, and recalling that DNIC are abolished by spinal atipamezole^4^, we investigated the impact of LC:LC or LC:SC module optoactivation on its expression. Interestingly, LC:SC opto-activation abolished DNIC, while LC:LC opto-activation only marginally decreased its potency (Fig. 2a-c). Meanwhile, while prazosin partially restored the light-evoked decrease in DNIC expression (Fig. 2d-g), atipamezole facilitated it (Fig. 2h-k). We hypothesise that the LC:SC circuit includes an inhibitory projection ‘branch', postsynaptic α_2_-AR mediated, to the DNIC origin nucleus. Upon opto-activation of the LC:LC module it is likely that a proportion of dorsal as well as ventral LC neurons are stimulated, explaining the decrease in DNIC potency. Thus, the dorsal LC is likely involved in governing DNIC functionality while the LC:SC direct pathway inhibits WDR activity via spinal α1-ARs. To investigate this separation of LC:SC and DNIC pathway functionality further we injected the neurotoxin DSP4 to deplete noradrenergic projections from the LC^11,12^ (Fig. 3a). No impact on WDR activity (Fig. 3b-d), was observed and DNIC were expressed (Fig. 3e). Elsewhere, we microinjected lidocaine (sodium channel blocker) to the LC ipsilateral to the recorded WDR neuron (Fig. 3f). This had no effect on WDR evoked activity (Fig. g-h), nor DNIC expression (Fig. 3i-j).

**Figure 1.**
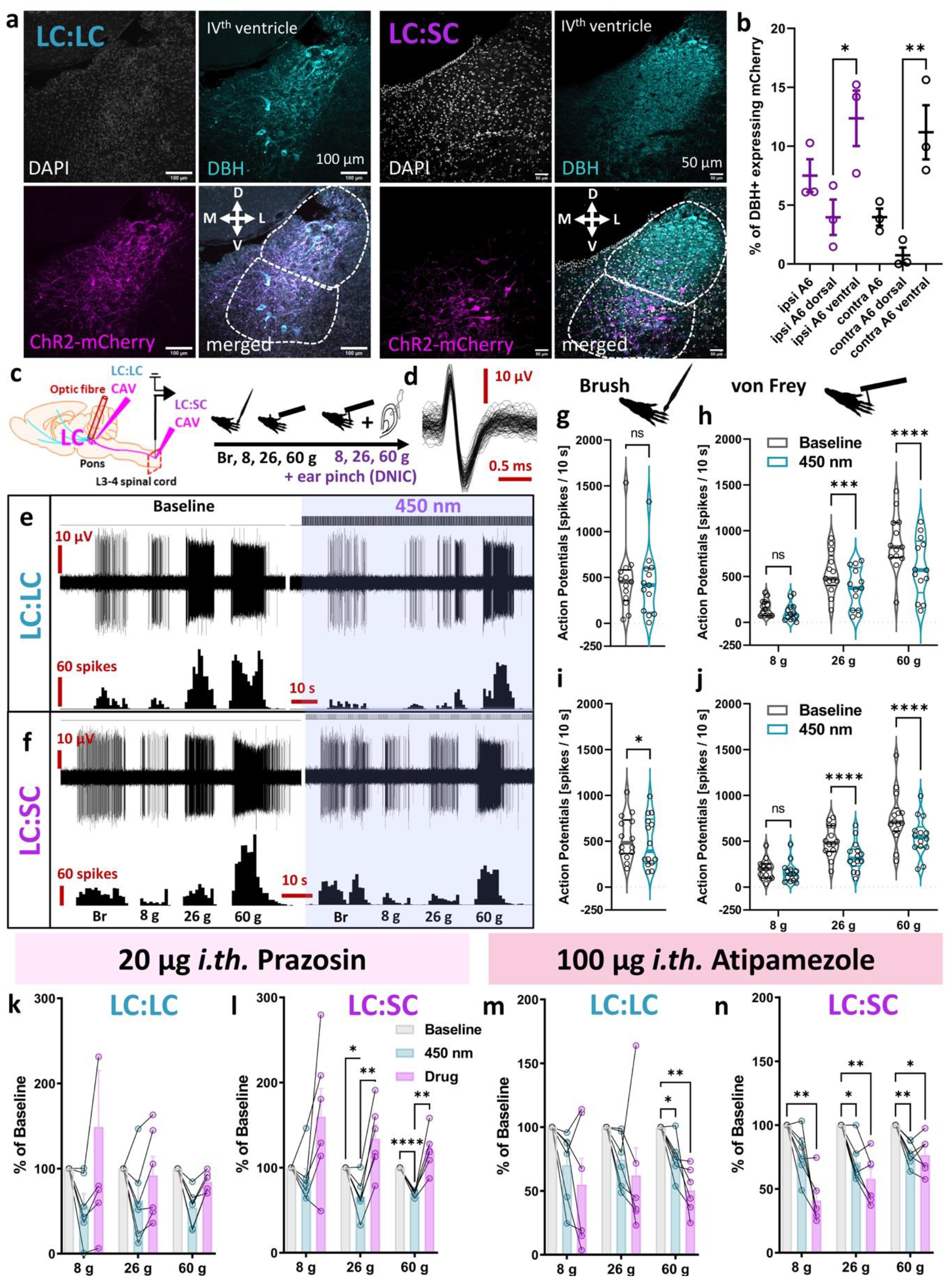
Heterogenous coerulean noradrenergic neuron population differentially modulate spinal nociception. **a)** Immunohistochemical analysis of locus coeruleus (LC) dopamine-β-hydroxylase (DBH)-expressing noradrenergic neurons transduced by canine adenovirus (CAV) delivering channelrhodopsin-2-mCherry construct under catecholamine specific promoter (PRS) injected locally (LC:LC module) or in the ipsilateral lumbar dorsal horns (LC:SC module). **b)** Percentage of mCherry-expressing DBH neurons in the ipsi- and contralateral LC following LC:SC module labelling. Mean±SEM of N=3 animals per group, n=6-8 slices per animal, unpaired One-Way ANOVA performed on N, [structure] P=0.0018, F_(5, 12)_=7.747. **c)** Schematic of the *in vivo* electrophysiological experiments. **d)** WDR neuron units code upon stimulation with natural stimuli. **e)** WDR neuron inhibition following LC:LC module ChR2-mediated activation (450 nm laser pulses: 5 Hz, 20 ms, 238 mW/mm^2^) **f)** The equivalent LC:SC module activation is shown. Quantification of **g)** brush and **h)** von Frey (vF) evoked activity before/after LC:LC module activation. **Brush and vF:** mean±SEM of N=13 animals per group, n=13 cells per group; Paired t-test performed on n: P>0.05 (brush) and Two-Way ANOVA (vF) performed on n, [vF] P<0.0001, F_(2, 36)_=24.37, [450 nm] P<0.0001, F_(1, 36)_=47.29. Quantification of **i)** brush and **j)** vF evoked activity before/after LC:SC module activation. **Brush and vF:** mean±SEM of N=14 animals per group, n=14 cells per group; Paired t-test performed on n: P<0.05 (brush); Two-Way ANOVA (vF) performed on n, [vF] P<0.0001, F_(2, 39)_=23.75, [450 nm] P<0.0001, F_(1, 39)_ = 89.83. Prazosin (α_1_-adrenoreceptors antagonist) reversed the inhibitory effect of **k)** LC:LC and **l)** LC:SC module activation. **LC:LC or LC:SC prazosin:** mean±SEM shown as percentage of baseline for N=6 animals per group, one cell per animal; Two-Way ANOVA with Geisser-Greenhouse correction [LC:LC-group] P>0.05, F_(1.03, 5.17)_=3.306 and [LC:SC-group] P<0.01, F_(1.23,6.17)_=14.24, respectively. **m)** LC:LC- and **n)** LC:SC-mediated inhibition of WDR neurons was further potentiated after spinal application of 100 μg atipamezole (α_2_-adrenoreceptors antagonist). **LC:LC or LC:SC atipamezole:** mean±SEM shown as percentage of baseline for N=6 animals per group, one cell per animal; Two-Way ANOVA with Geisser-Greenhouse correction [LC:LC-group] P<0.05, F_(1.39, 6.94)_=5.635, and [LC:SC-group] P<0.001, F_(1.82, 9.09)_=26.58. Tuckey *post-hoc* used for all ANOVA: *P<0.05, **P<0.01, ***P<0.001, ****P<0.0001. See also Supplementary figure 1.

**Figure 2.**
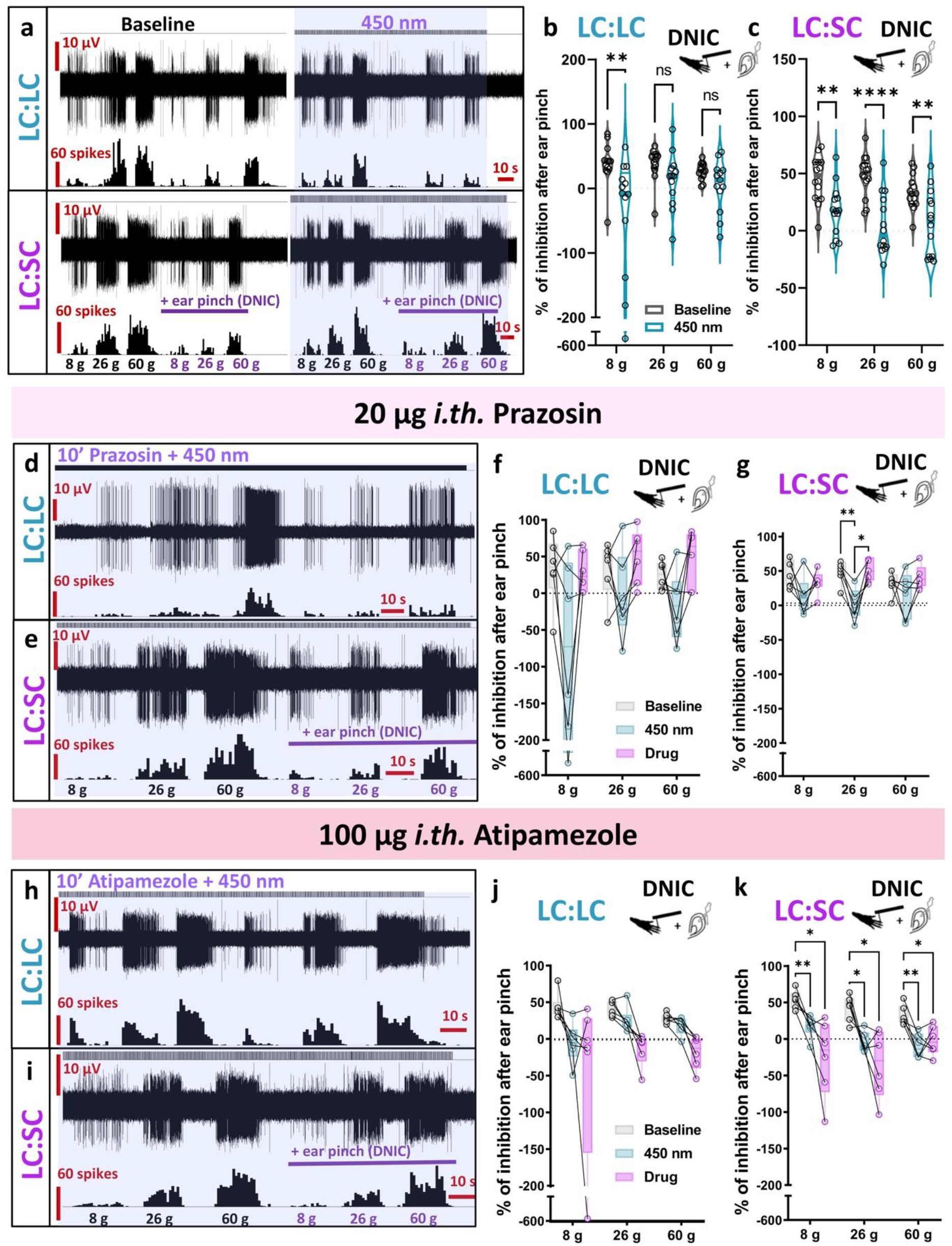
Differential modulation of diffuse noxious inhibitory controls (DNIC) expression by coerulean neurons. **a)** DNIC expression, quantified as the inhibitory effect of a conditioning stimulus (ear pinch), decreased following LC:LC module optoactivation (450 nm laser pulses), and was abolished following identical LC:SC module activation. **b)** Percentage of inhibition after DNIC activation before/after LC:LC module activation. Mean±SEM of N=13 animals per group, n=13 cells per group; Two-Way ANOVA performed on n, [450 nm] P<0.01, F_(1, 36)_=10.75. **c)** Identical experiments before/after the LC:SC module activation. Mean±SEM of N=14 animals per group, n=14 cells per group; Two-Way ANOVA performed on n, [450 nm] P<0.0001, F_(1, 39)_=46.01. Prazosin partially reversed the impact of **d)** LC:LC or **e)** LC:SC module activation on DNIC expression: **f)** LC:LC prazosin: mean±SEM shown as percentage of baseline for N=6 animals per group, one cell per animal; Two-Way ANOVA with Geisser-Greenhouse correction [group] P<0.05, F_(1.38, 6.91)_=8.056. **g) LC:SC prazosin:** mean±SEM shown as percentage of baseline for N=6 animals per group, one cell per animal; Two-Way ANOVA with Geisser-Greenhouse correction [group] P<0.05, F_(1.17, 5.87)_=8.215. The inhibitory effect of **h)** LC:LC and **i)** LC:SC module activation on DNIC expression was facilitated by spinal atipamezole: **j) LC:LC atipamezole:** mean±SEM shown as percentage of baseline for N=6 animals per group, one cell per animal; Two-Way ANOVA with Geisser-Greenhouse correction [group] P>0.05, F_(1.06, 5.31)_=4.950. **k) LC:SC atipamezole:** mean ± SEM shown as percentage of baseline for N=6 animals per group, one cell per animal; Two-Way ANOVA with Geisser-Greenhouse correction [group] P<0.001, F_(1.82, 9.01)_=26.58. Tuckey *post-hoc* used for all ANOVA: *P<0.05, **P<0.01, ****P<0.0001.

**Figure 3.**
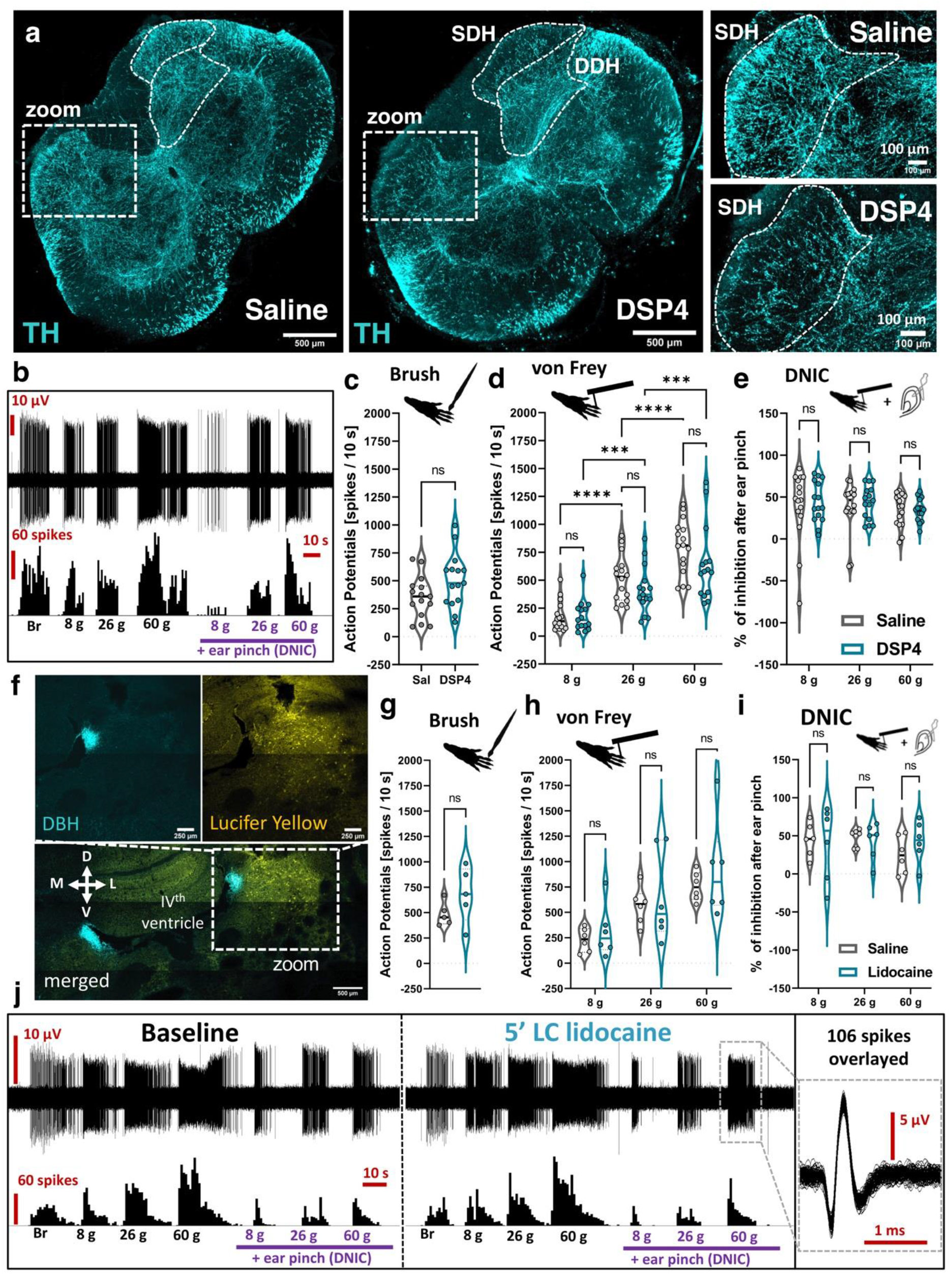
Ablation of coerulean noradrenergic fibres does not affect basal spinal convergent neuron activity nor diffuse noxious inhibitory controls (DNIC) expression. **a)** A PACT-cleared 500 μm thick lumbar spinal cord section (saline versus DSP4-treated rats) evidences a decrease in tyrosine hydroxylase (TH) immunolabelled fibres in the superficial but not deep dorsal horn (SDH/DDH). **b)** DSP4 treatment did not impact WDR neuron sensory coding nor DNIC expression. Quantification of **c)** brush and **d)** von Frey (vF) evoked action potentials in saline and DSP4 treated rats. **Brush and vF:** mean±SEM of N=6 animals per group, n=15 cells per group; unpaired t-test performed on n: P>0.05 (brush); Two-Way ANOVA (vF) performed on n, [vF] P<0.0001, F_(2, 30)_=128.7, [DSP4] P>0.05, F_(1, 15)_=1.851. **e)** Percentage of inhibition after DNIC activation as shown in b). **DNIC:** mean±SEM of N=6 animals per group, n=15 cells per group; Two-Way ANOVA performed on n, [DSP4] P>0.05, F_(1, 15)_=0.2105. **f)** Analogously to DSP4 treatment, ipsilateral LC microinjection of 2% lidocaine (marked by Lucifer Yellow) does not affect WDR neuronal activity nor DNIC expression, as quantified in **g)** for brush and **h)** for vF. **Brush:** mean±SEM of N=5 animals per group, n=5 cells per group; paired t-test performed on n: P>0.05. **vF:** mean±SEM of N=6 animals per group, n=6 cells per group; Two-Way ANOVA performed on n, [vF] P<0.001, F_(2, 10)_=23.97, [DSP4] P>0.05, F_(1, 5)_=0.78. **i)** Percentage of inhibition after DNIC activation as shown in **j)**. **DNIC:** mean±SEM of N=6 animals per group, n=6 cells per group; Two-Way ANOVA performed on n, [DSP4] P>0.05, F_(1, 5)_= 0.063. Tuckey *post-hoc* used for all ANOVA: *P<0.05, **P<0.01, ****P<0.0001.

It is likely that maladaptive reciprocal communication between the LC and DNIC-origin nuclei underlies certain chronic pain phenotypes. Previous research has demonstrated abolished DNIC expression in the late stage of a model of chronic joint inflammatory pain and impaired descending noradrenergic modulation with relation to the LC^13^. This insight, specifically linking stage-specific DNIC attenuation to impaired LC functionality, lends weight to the argument that reciprocity between LC and DNIC origin nuclei governs the final output of descending modulatory controls that are subserved by noradrenaline. However, the nature of the influence is unknown.

The coerulean neuronal population is developmentally diverse^14^ with distinct anatomical projections^5^. Output from the ventral LC proffers an analgesic spinal α1-ARs-mediated pathway, whereas output from the dorsal LC is proalgesic^5^. The underlying mechanism(s) will include neuro-immune interactions since superficial dorsal horn astrocytes expressing α_1_-ARs were shown to be critical analgesic regulators in monoaminergic transmission terms^15^. Crucially, LC and DNIC projection pathways offer distinct mechanisms of naturally occurring pain-relief. Thus, ‘one size doesn't fit all' when adopting a therapeutic approach to reinstate DPMS activity because there are multiple descending inhibitory pathways therein that are evolutionarily distinct despite overlapping functionality.

## Acknowledgements

Authors would like to thank Professor Anthony Pickering for supplying us with CAV virus used in this study and Professor Stephen McMahon for start-up equipment funding.

---

**Supplementary figure 1.**
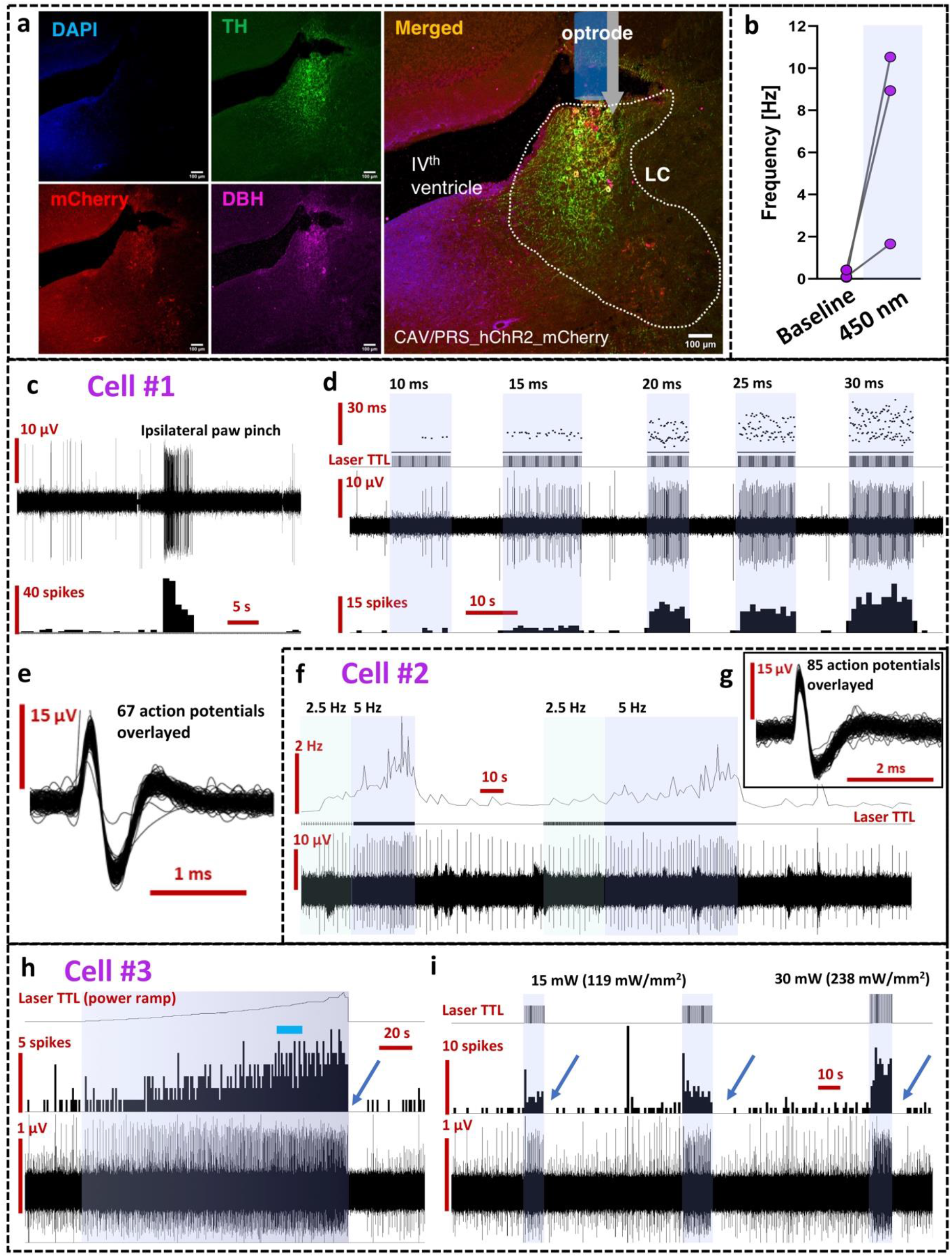
Optimisation of light parameters for locus coeruleus (LC) neuron optoactivation. **a)** A representative coronal section of transduced LC after in situ injection of CAV/PRS-hChR2-mCherry. Tyrosine hydroxylase (TH) and dopamine-β-hydroxylase (DBH) immunolabelled cells overlapped with the virally-delivered mCherry in the LC proper. **b)** Mean firing frequency of transduced LC before and after optoactivation with chosen optimal 450 nm laser light parameters: 30 mW (238 mW/mm^2^), 20 ms pulse, at 5 Hz. **c)** LC neuron responding to ipsilateral noxious pinch. Basal spontaneous discharges are shown before and after pinch. Note around 15 s inactivity period immediately after pinch. **d)** Same neuron as in b) optoactivated with different light pulse duration, but same frequency (5 Hz) and power (238 mW/ mm^2^). 20 ms pulse width was chosen as optimal. **e)** Overlayed action potentials of cell shown in c) and d). **f)** Optoactivation of the LC cell from different animal. Matching power (238 mW/ mm^2^) and pulse width (20 ms) was used, but with two different frequencies (2.5 and 5 Hz). 5 Hz was chosen optimal. **g)** An inclusion showing overlayed action potentials of the cell recorded in f). **h)** Light power ramp performed on another optically-sensitive LC neuron. An increase of discharges with the increase of laser power is shown. Blue bar indicates chosen power (238 mW/ mm^2^). Note a 15 s inactivity period immediately after laser light was turned off (blue arrow). **i)** Same cell as in h) showing light evoked responses with two different laser powers, but similar temporal pattern of pulses (20 ms width at 5 Hz).

## Materials and Methods

### Animals

Male Sprague-Dawley rats (Envigo, UK) were used for experiments. Animals were group housed on a 12:12 h light–dark cycle. Food and water were available *ad libitum.* Animal house conditions were strictly controlled, maintaining stable levels of humidity (40–50%) and temperature (22 ± 2°C). All procedures described were approved by the Home Office and adhered to the Animals (Scientific Procedures) Act 1986. Every effort was made to reduce animal suffering and the number of animals used was in accordance with International Association for Study of Pain (IASP)^1^ and ARRIVE ethical guidelines^2^.

The observer was blind to DSP4 and LC:LC/LC:SC labelled experimental groups, but not to the pharmacological treatment performed during electrophysiological recordings. All surgical procedures were carried out under aseptic conditions and animals were maintained at 37°C on a homeothermic pad. Brainstem tissue from injected animals was histologically analysed *post hoc* to confirm transduction of LC neurons (fluorescent tag expression).

All experiments were designed to contain minimum of 6 rats per group, based on G-power predictions from previous experiments. Animals were randomly assigned to experimental groups. From 60 rats designated for this study, 7 failed to provide stable WDR neuronal recordings, 3 rats developed vestibular problems reaching humane endpoint within 4 days after LC virus microinjection, and 1 animal died 24 hours after DSP4 administration. In total 49 rats were used as follows: 24 rats were used for mixed opto-pharmacology experiments (6 rats per group: LC:LC atipamezole, LC:LC prazosin, LC:SC atipamezole, LC:SC prazosin). Additional 3 rats were used for WDR baseline characterisation with optogenetics (2 for LC:SC, 1 for LC:LC) followed by LC optoelectrical recordings of transduced neurons. In the latter no pharmacology was performed. Further 6 rats were used for DSP4 group and 15 rats were used as naïve controls, which included 6 rats used for lidocaine microinjection experiment.

### DSP4 injections

For ablation of the coerulean noradrenergic fibres across the neuroaxis, a selective neurotoxin N-(2-chloroethyl)-N-ethyl-2-bromobenzylamine (DSP4) hydrochloride (Sigma, Dorset, UK) was used ^3,4^. Six rats weighting 60-80 g were injected intraperitoneally with freshly prepared 50 mg/kg DSP4 in saline. Additional six control rats received vehicle injection. 15% lethality rate from the DSP4 toxin is expected^5^, therefore, to minimise eventual animals suffering injections were given early on the day and rats were kept for up to 4 hours in the recovery incubator (set at 32°C) and carefully monitored for up 8 hours to minimise lethality. 14-16 days after the toxin injection terminal electrophysiological characterisation of lumbar deep dorsal horn WDR neurons was performed followed by transcardial perfusion with cold saline followed by 4% PFA and tissue collection for histological analysis.

### Virus injections

#### The LC:LC module

To transduce catecholaminergic coerulean neurons, ipsilateral LC stereotaxic injections were made (Kopf Instruments, UK) analogously to described in detail earlier^6^. In brief, male Sprague-Dawley rats (180-220 g, Charles River) were anesthetized with a mix of ketamine (5 mg/100 g, Vetalar; Pharmacia) and medetomidine (30 μg/100 g, Dormitor; Pfizer) delivered intraperitoneally until loss of paw withdrawal reflex and perioperative analgesia was achieved by the subcutaneous injections of meloxicam (2 mg/kg, Metacam^®^, Boehringer Ingelheim, Berkshire, UK). The animal was placed in a stereotaxic frame and core temperature was maintained at 37°C using a homeothermic blanket (Harvard Apparatus, US). Aseptic surgical techniques were used throughout, and eyes were protected with sterile paraffin-based moisturiser (Lacri-Lube, Allergan, UK). Using a 0.7 mm dental drill a hole was made in the skull right above the targeted structure and dura was carefully pierced with the aid of surgical microscope until the visible amount of CSF appeared. With 10° rostral angulation to avoid puncturing the sinus, the following coordinates were used (from lambda): RC: −2.1 mm, ML: 1.3 mm, and −5.8-6.2 mm deep from the cerebellar surface. A glass pulled micropipette coupled to electronically controlled nanoinjector (Nanoliter 2010, WPI, FL, US) with back-filled mineral oil as a medium was used for precise delivery of canine adenovirus (CAV) carrying channelrhodopsin 2 under the control of catecholamine-specific synthetic promoter (sPRS) (CAV-sPRS-hChR2(H134R)-mCherry, titer >3×10^10^ TU/ml, PVM, Montpellier, a gift from Professor Anthony Pickering, University of Bristol ^6,7^). The micropipette maintaining 20-40 μm tip diameter was front loaded with the virus immediately prior the injection. Three injections, each of 400 nl, were made every 200 μm starting from −6.2 mm (DV) with 2 nl/s delivery rate and minimal 3-5 minutes were allowed between slow pipette retraction. The wound was irrigated with saline and closed with Vicryl 4-0 absorbable sutures and wound glue (VetBond, 3M, UK). Anaesthesia was reversed with i.p. injection of atipamezole (Antisedan, 0.1 mg/100 g; Pfizer, UK). The animals were placed in a thermoregulated recovery box until fully awake. Two to three weeks were allowed for the transgene expression, after which animals were taken for terminal *in vivo* electrophysiology.

#### The LC:SC module

To transduce spinally projecting catecholaminergic brainstem neurons, the same CAV-sPRS-hChR2(H134R)-mCherry virus (titer >3×10^10^ TU/ml, ^7^) was injected in the lumbar spinal cord. In brief, male Sprague-Dawley rats weighting 60-80 g were used for injections. Following the induction of anaesthesia (5% isoflurane in 1 l/min oxygen) rats were placed in a stereotaxic frame (without fixing the head) and maintained with 1.8-2% isoflurane in oxygen (1 l/min flow) delivered via nose cone. Core temperature was maintained at 37°C using a homeothermic blanket (Harvard Apparatus, US), aseptic surgical techniques were used throughout and eyes were protected with sterile paraffin-based moisturiser (Lacri-Lube, Allergan, UK). The subcutaneous injections of meloxicam (2 mg/kg, Metacam, Boehringer Ingelheim, Berkshire, UK) were used for perioperative analgesia. After a complete loss of paw withdrawal reflex the 2-2.5 cm skin overlaying lumbar spinal cord was incised and overlaying muscles carefully dissected to gain access to T12/L1 intervertebral space. The sterile spinal clamp was used to secure the cord and the spinal L3-L4 region was exposed by bending the cord rostrally providing easy access to the underlaying dura without the need for extensive laminectomy. A small puncture in the dura was made with the aid of surgical microscope and a glass pulled micropipette (20-40 μm tip diameter) was inserted in the dorsal horn to carefully deliver CAV virus at 2 nl/s with the aid of electronically controlled nanoinjector (Nanoliter 2010, WPI, FL, US). Three injections, each of 400 nl, were made ipsilaterally at around 300-400 μm apart in the rostrocaudal axis, 200 μm lateral from the central vessel, and at 800, 600 and 400 μm deep from the cord surface for injection 1-3, respectively. After each injection minimum of 5 minutes were allowed for the virus diffusion in the cord parenchyma followed by slow pipette retraction. The surgical area was maintained moist with the aid of sterile saline. The wound was closed with sterile wound clips (removed at day 7 post-surgery) and surgical glue (VetBond 3M, UK). The animals were placed in a thermoregulated recovery box until fully awake. Two to three weeks were allowed for the transgene expression, after which animals were taken for terminal *in vivo* electrophysiology.

#### Spinal Cord In Vivo Electrophysiology

*In vivo* electrophysiology was performed on animals weighing 240–300 g as previously described ^8^. Briefly, after the induction of anaesthesia, a tracheotomy was performed, and the rat was maintained with 1.5% of isoflurane in a gaseous mix of N_2_O (66%) and O_2_ (33%). Core body temperature was monitored and maintained at 37 °C by a heating blanket unit with differential rectal probe system. Electrocardiogram (ECG) was monitored by two intradermal needles inserted in front limbs with signal amplified by the Neurolog system consisting of AC preamplifier (Neurolog NL104, gain x200), through filters (NL125, bandwidth 300 Hz to 5 KHz) and a second-stage amplifier (Neurolog NL106, variable gain 600 to 800) to an analogue-to-digital converter (Power 1401 625kHz, CED). Craniotomy was performed to gain stereotaxic access to the ipsilateral LC for either optic fibre or micropipette insertion as described in following sections. A laminectomy was performed to expose the L3–L5 segments of the spinal cord, the cord was clamped to minimise movement, dura was carefully removed with the aid of surgical microscope, and the recording area was secured by saline-filled well made in solidified 2% low melting point agarose (made in saline also). Using a parylene-coated, tungsten electrode (125 μm diameter, 2 MΩ impedance, A-M Systems, Sequim, WA, USA), wide dynamic range neurons in deep laminae IV/V (~650–900 μm from the dorsal surface of the cord) receiving intensity-coding afferent A-fibre and C-fibre input from the hind paw were sought by periodic light tapping of the glabrous surface of the hind paw. Extracellular recordings made from single neurones were visualized on an oscilloscope and discriminated on a spike amplitude and waveform basis. Specifically, the signal from the electrode's tip was processed via headstage connected to the neurolog system consisting of AC preamplifier (Neurolog NL104, gain x200), through HumBag (Quest Scientific, North Vancouver, Canada) used to remove low frequency noise (50–60 Hz), via a second-stage amplifier (Neurolog NL106, variable gain 600 to 800), filters (NL125, bandwidth 1000 Hz to 5 KHz) and spiketrigger (Neurolog NL106, variable gain 600 to 800) to an analogue-to-digital converter (Power 1401 625kHz, CED). Spike trigger was visualised on a second oscilloscope channel and manually set to follow single unit spikes. Its analogue signal was digitalised via event input to build stimulus histogram in real time along the waveform recordings. All the data were captured by an analogue-to-digital converter (Power 1401 625kHz, CED) connected to a PC running Spike2 v8.02 software (Cambridge Electronic Design, Cambridge, UK)) for data acquisition, analysis and storage. Simultaneous ECG monitoring and transistor–transistor logic (TTL) triggers (i.e. for the lasers, see below) were additionally coupled to Spike 2 recording traces via CED-1401 analogue inputs.

#### Stimulation paradigm in all electrophysiological recordings

Natural mechanical stimuli, including brush and von Frey filaments (8 g, 26 g and 60 g) and von Frey filaments with concurrent ipsilateral noxious ear pinch (15.75 × 2.3 mm Bulldog Serrefine ear clip; InterFocus, Linton, United Kingdom), were applied in this order to the receptive field for 10 s per stimulus. The noxious ear pinch was used as a conditioning stimulus (CS) to trigger diffuse noxious inhibitory control (DNIC, ^9^) and was quantified as an inhibitory effect on neuronal firing during ear pinch to its immediate respective von Frey filament applied without the conditioning stimulus (% of inhibition after ear pinch). A minimum 30 s nonstimulation recovery period was allowed between each test in the trial. A 10-minute nonstimulation recovery period was allowed before the entire process was repeated for control trial number 2 and 3. The procedure was repeated 3 times and averaged only when all neurons met the inclusion criteria of 10% variation in action potential firing for all mechanically evoked neuronal responses. No animals were excluded from analysis. Spontaneous discharges of spinal WDR neurons were quantified *post hoc* as an averaged discharge frequency in 300 s (5 minutes) periods of inactivity. These periods were immediately preceding stimulation trials and at least 300 s after the last stimulation period.

#### In vivo spinal pharmacology with electrophysiological monitoring

After collection of predrug baseline control data as outlined above, atipamezole (a α_2_-AR antagonist: 10 and 100 μg; Sigma-Aldrich, Gillingham, United Kingdom, dissolved in 97% normal saline, 2% Cremophor [Sigma, UK], 1% dimethyl sulfoxide [DMSO; Sigma, UK] vehicle), prazosin hydrochloride (α_1_-AR antagonist: 20 μg, Sigma-Aldrich, Gillingham, United Kingdom, dissolved in water for injections) was administered topically to the spinal cord in 50 μl volumes following gentle removal of residing saline in the agarose well. Each individual drug dose effect (one stable neuron assessed per rat) was followed for up to 60 minutes with tests performed typically at 3 time points (starting at 10, 20 and 30 minutes). For each time point, a trial consisted of consecutive stable responses to brush, von Frey and DNIC (von Frey with concurrent ipsilateral ear pinch). The post-drug effects in subsequent individual modalities (as compared to mean pre-drug baseline) were judged by the maximal change in recorded action potential rate. All data plotted represents the time point of peak change based on these criteria.

#### Lidocaine block of the LC activity

Six naïve rats weighting 240-260 g were used for lidocaine block of neuronal activity in the LC with simultaneous terminal spinal WDR neuron electrophysiological recordings. In brief, a hole in the skull was drilled right prior the lumbar laminectomy for DDH WDR neuronal recordings as described above. Utilising the same parameters to reach the ipsilateral to the spinal recording side ventral LC (−6.0 mm from the cerebellum surface), injections of lidocaine (2% in saline, Cat. L5647 Sigma, UK) were made after collecting stable baseline recordings of DDH WDR neurons. A glass pulled pipette was used coupled to the electronically controlled nanoinjector (Nanoliter 2010, WPI, FL, US) to precisely deliver the drug (500 nl, 2 nl/s). After the drug delivery, pipette was left in place throughout the spinal recordings. Tests were performed at 5-, 10- and 15-minutes post lidocaine injection. At the end of the experiment the solution in the pipette was replaced with 0.5% Lucifer Yellow-CH dipotassium salt (#L0144, Sigma, UK) and the pipette was re-inserted in the same brain region and 500 nl of the Lucifer Yellow Solution was injected therein. 10 minutes were allowed for the dye diffusion after which the animal was sacrificed by the anaesthetic overdose and immediate transcardial perfusion followed by brain extraction for anatomical verification post-mortem.

### Optogenetics

#### Light stimulation of coerulean neurons during spinal WDR recordings

The 450 nm laser (Doric Lenses, Quebec, Canada) was externally TTL-triggered by the neurolog system (NeuroLog system, Digitimer, UK) to deliver defined light pulses (20 ms pulse width at 5 Hz). In brief, the laser light was coupled to a multimode 200 um patch cord (0.39 NA, #M75L01, Thorlabs, UK) and via SMA to SMA mating sleeve (#ADASMA, Thorlabs, UK) to second ferrule-terminating multimode 200 μm patch cord (0.39 NA, #M77L01, Thorlabs, UK) directly interconnected to multimode stainless steel 20 mm long cannula (200 μm diameter, 0.39 NA, #CFM12L20, Thorlabs, UK). The laser controller current was set to 44-54 mA adjusted (using PM16-130 power meter, Thorlabs, UK) to correspond to 30 mW (238 mW/mm^2^) light power density at the tip of the implantable 200 μm fibre cannula ^6^. The power density was adjusted for each preparation. After desired power was achieved the cannula was slowly inserted in the ventral LC ipsilateral to the recorded spinal WDR neurons. The canula was lowered using precise hydraulic micromanipulator (Narishige, Japan) with 10° rostral angulation at the following coordinates (from lambda): RC: −2.1 mm, ML: 1.3 mm, and −6.0 mm deep from the cerebellar surface.

Spinal WDR neurons were characterised by three stable baseline responses followed by three optically modulated responses. For combined optogenetics and spinal pharmacology, after collecting three stable baseline and three stable optoactivation responses (averaged, if stable), a drug (either 100 μg atipamezole or 20 μg prazosin) was applied topically on the exposed spinal cord surface, right above the recording site. To assess simultaneous action of the drug and the LC optoactivation, light pulses were delivered 30 s before and throughout each series of tests (approximately 5 minutes per series) and minimally 5 minutes of the recovery time was allowed between the tests. Pharmacology was monitored every 10 minutes for 30-40 minutes (each test with optoactivation) and the 60 minutes time point was to test neuron returning to the baseline (no optoactivation). At the end of every experiment, animals were sacrificed by the overdose of isoflurane and transcardially perfused with cold saline followed by 4% paraformaldehyde for anatomical evaluation.

#### LC neuron recording and optoactivation

A simultaneous recording and optical stimulation of the transduced LC neurons were made using microoptrodes as described earlier with minor modifications ^10^. LC neurons were identified as described before ^6^ by their large amplitude with duration of action potentials over 1 ms, spontaneous firing (0.5-7 Hz) and biphasic response following hindpaw pinch (activation followed by transient silent period, particularly strong for the contralateral hindpaw). For the LC recordings and optoactivation, the all-glass recording microptrode with 20 μm tip diameter consisted of the recording core filled with 3 M sodium acetate (resulting in 2-3 MΩ resistance) and the parallel gradient index (GRIN) optical core (a gift from Professor Yves De Koninck, Laval University, Canada) coupled to the optic fibre (multimodal, 200 μm core diameter, 0.39 NA, #M77L01, Thorlabs, UK) was used. The LC neurons were optoactivated by 450 nm laser (Doric Lenses, Quebec, Canada) light pulses as described above ^6^. With constant laser power set as 30 mW (238 mW/mm^2^) and 5 Hz frequency we tested 10, 15, 20, 25 and 30 ms wide pulses, choosing 20 ms as the optimal pulse based on its almost maximal activation.

#### Immunohistochemistry

Animals were sacrificed by the overdose of anaesthetic and transcardially perfused with cold phosphate buffer saline (PBS) followed by 4% paraformaldehyde (PFA) in phosphate buffer (pH 7.5). Next, collected spinal cords and brains were post-fixed in 4% PFA for 3-4 days at 4°C, followed by 3-4 days at 4°C in 30% sucrose. Once tissue density equilibrated, lumbar spinal cords and brains were precut into 5 mm thick coronal fragments with razor blades and the aid of rat brain matrix. Obtained fragments transferred to optimum cutting temperature (OCT)-filled moulds were snap-frozen in liquid nitrogen and stored frozen until further analysis. Next the OTC embedded tissue was cryo-sectioned (Bright Instruments, UK) to 25 μm thick coronal slices subsequently collected on eight Menzel-Gläser Superfrost Plus slides (a slice collected every 200 μm) and stored in −20°C until staining. Once dried (45°C for an hour) and briefly washed with 50% ethanol, sections were outlined with a hydrophobic marker (PAP pen, Japan), rehydrated and blocked with 10% donkey serum in blocking solution (0.03% NaN_3_, 0.3% Triton X-100 in PBS, pH=7.5) for two hours prior to overnight incubation at room temperature with primary antibodies against dopamine-β-hydroxylase (DBH, a marker of noradrenergic neurons: Mouse, 1:500, Millipore, MAB308, UK), mCherry (Rabbit, 1:500, Abcam, ab167453, UK). Slides were then PBS washed and incubated with the appropriate fluorophore-conjugated secondary antibodies in blocking solution (Donkey anti-Rabbit, Alexa Fluor 568, A10042, Invitrogen, Eugene, OR, US; Donkey anti-Mouse, AlexaFluor 488, A21202, Invitrogen, Eugene, OR, US; all used at 1:1000 dilution) for 4 hours to overnight at room temperature. Slides were protected with mounting media (Fluoromount-G with DAPI, eBioscience, UK) and coverslips and stored in darkness at 4°C until imaging.

Samples were typically imaged with an LSM 710 laser-scanning confocal microscope (Zeiss) using Zeiss Plan Achromat 10x (0.3 NA) and 20 x (0.8 NA) dry objectives and analysed with Fiji Win 64. For quantification, samples were imaged with 20x dry objective on Zeiss Imager Z1 microscope coupled with AxioCam MRm CCD camera. The acquisition of images was made in multidimensional mode and the MosaiX function was used to construct the full view. 6-8 slices were imaged per animal. Cell counting was carried out on the Fiji Win 64 utilising cell counter plugin. On average, 20-30 brainstem sections were imaged for quantification.

#### Passive Tissue Clearing (PACT)

A passive CLARITY tissue clearing technique (PACT) (described in detailed in: ^11^) has been implemented to allow thick (1000-2500 μm) spinal cord fragments imaging. Briefly, following transcardial perfusion of deeply anaesthetised rats with a cold PBS and a cold 4% PFA in phosphate buffer, pH=7.5, spinal cords were extracted and post-fixed in 4% PFA for 3-4 days in 4°C. After fixation, samples were transferred directly to ice-cold A4P0 solution consisting of: 4% acrylamide monomer (40% acrylamide solution, cat. 161-0140, Bio-Rad, UK), 0.25% VA-044 (thermoinitiator, Wako, US) in 0.01 M PBS, pH=7.4, and incubated at 4°C overnight in distilled water prewashed (to remove anticoagulant) vacutainer tubes (Vacutainer, #454087, Greiner GmbH, Austria). The next day, samples were degassed by piercing the septum with a 20G needle connected to a custom-build vacuum line. The residual oxygen was replaced with nitrogen by 2 min bubbling of the solution with pure nitrogen (BOC, UK) via a long, bottomreaching 20G needle, and a second short needle pierced to allow gases to exhaust. Throughout, samples were kept on ice to prevent heating and consequent premature A4P0 polymerisation. After achieving oxygen-free conditions, samples were polymerised by 3 h incubation in a 37°C water bath. Following polymerisation, the excess honey-like polyacrylamide gel was removed with tissue paper, and samples were transferred to 50 ml falcon tubes filled with clearing solution. 10% SDS (#L3771, Sigma-Aldrich, UK) in PBS, pH=8.0, was used for passive clearing. Samples were incubated on a rotary shaker at 37°C and 75 rpm (Phoenix Instruments, UK) until reaching the appropriate transparency (usually 3-4 days).

Next, all samples were washed extensively with PBS, pH=7.5, on rotary shaker at room temperature, by replacing the solution 4-5 times throughout the course of 1 day in order to remove the SDS. Following washing, samples were treated with primary antibody in blocking buffer consisting of 2% normal donkey serum in 0.1% Triton X-100 in PBS, pH=7.5 with 0.03% sodium azide. 400 μl of rabbit anti-tyrosine hydroxylase (TH, marker of catecholaminergic neurons; 1:250, Ab152, Millipore, UK) primary antibody was used per 2 mm thick spinal cord slice in a 2 ml Eppendorf tube. Samples were incubated with the primary antibody at room temperature, with gentle shaking for 3 days. This was followed by 4-5 washing steps with PBS over the course of a day days. Next, the samples were incubated with gentle agitation, at room temperature, in darkness, for 3 days with the goat anti-rabbit (Alexa Fluor 647, A21244, Invitrogen, Eugene, OR, US) fluorophore-conjugated secondary antibody (1:200 in the blocking buffer). Thereafter, samples were washed extensively with PBS at least 5 times over 1-2 days at room temperature. Finally, samples were overnight incubated in the refractive index-matching solution (RIMS, refractive index = 1.47) consisting of 40 g of Histodenz (#D2158, Sigma-Aldrich, UK) dissolved in 30 ml of PBS, pH=7.5 with 0.03% sodium azide. 400-600 μl of RIMS was used per structure. Samples were placed in fresh RIMS in custom-made glass slide chambers, covered with coverslips, and equilibrated for few hours before imaging.

Samples were imaged with a Zeiss LSM 780 one-photon confocal upright microscope, equipped with EC Plan-Neofluar 10x 0.3 NA, Ph1 dry objective (WD=5.3 mm, cat. 420341-9911, Zeiss, Germany) and 633 nm laser lines. Scans were taken with 2048×2048 pixel resolution, with 4-5 μm optical section typically spanning 400-700 μm of scanned depth (resulting in 100-150 planes) with auto Z-brightness correction to ensure uniform signal intensity throughout the sample. Images were exported from Zen 2012 Blue Edition software (Carl Zeiss Microscopy GmbH, Germany). Next graphical representations, 3D-rendering, animations, maximal intensity projections within selected z-stacks and further analysis were obtained with open-source Fiji (ImageJ) equipped with appropriate plugins.

#### Quantification and Statistical Analysis

Statistical analyses were performed using SPSS v25 (IBM, Armonk, NY, USA). All data plotted in represent mean ± SEM. Typically, up to 4 WDR neurons were characterised per preparation (n), and data were collected from at least 6 rats per group (N). Single pharmacological investigation was performed on one neuron per animal. Statistical analysis was performed either on number of neurons (n) for populational studies or number of animals (N) for pharmacological studies. Therefore, throughout the manuscript “n” refers to the number of cells tested and “N” to the number of animals tested. Detailed description of the number of samples analysed and their meanings, together with values obtained from statistical tests, can be found in each figure legend. Symbols denoting statistically significant differences were also explained in each figure legend. Main effects from analysis of variance (ANOVA) are expressed as an F-statistic and *p*-value within brackets. Throughout, a p-value below 0.05 was considered significant. One-way ANOVA with Tukey post-hoc test was used to assess significance for labelling efficiency between structures. Uncorrected two-way repeated-measures (RM) ANOVA with the Tukey post-hoc was used to assess von Frey and DNIC responses in the baseline conditions. For pharmacological experiments, Geisser-Greenhouse correction was used for RM-ANOVA. Paired student t-test was used to assess brush-evoked responses. GraphPad Prism was used to analyse the data.

